# Extracellular Matrix Reorganization During Endometrial Decidualization

**DOI:** 10.1101/2025.03.22.644728

**Authors:** Mona Gebril, Sparhawk Mulder, Rimi Das, Shanmugasundaram Nallasamy

**Affiliations:** Department of Obstetrics, Gynecology, and Reproductive Sciences, Larner College of Medicine University of Vermont, Burlington, VT

**Keywords:** Mouse Endometrium, Embryo Implantation, Decidualization, Extracellular Matrix, Collagen, Elastic Fibers, Lysyl Oxidases

## Abstract

Extracellular matrix reorganization, a concurrent process of endometrial decidualization, has garnered widespread recognition. However, our understanding of this process remains limited. In this study, we aimed to investigate the expression, spatial distribution, and reorganization of fibrillar collagens, elastin, and lysyl oxidases within the decidua. Using second harmonic generation imaging, we successfully recorded fibrillar collagen reorganization between preimplantation and decidualized endometrium. Upon embryo implantation, the fibrillar collagens align themselves parallel to the direction of embryo invasion. Furthermore, we employed confocal imaging analysis to reveal distinct expression and spatial distribution patterns of elastin and lysyl oxidase-like enzymes. Elastin expression begins to manifest surrounding the implanting embryo, extends into the decidua, and exhibits a high concentration in the mesometrial region after gestation day 8. All lysyl oxidase-like enzymes are localized within the decidua, although they exhibit varying expression patterns. To gain further insights, we utilized an *in vitro* stromal cell decidualization model and provided compelling evidence that stromal cells serve as the primary source of the extracellular matrix components during endometrial decidualization. Additionally, we demonstrated that the genes encoding factors involved in the synthesis, processing, and assembly of fibrillar collagen and elastic fibers exhibit differential expression patterns during *in vitro* decidualization. Genes such as asporin, decorin, thrombospondin 2, fibulin 2, fibulin 5, and lysyl oxidase show significant induction during *in vitro* decidualization. In summary, our comprehensive analysis provides a detailed evaluation of the expression, spatial distribution, and reorganization of fibrillar collagens, elastin, and lysyl oxidases during the process of endometrial decidualization.

**Summary Sentence:** Fibrillar collagen, elastic fibers, and lysyl oxidases, synthesized by endometrial stromal cells, exhibit distinct spatial distributions and reorganization patterns within the decidua during embryo implantation.

**Graphical Abstract:** 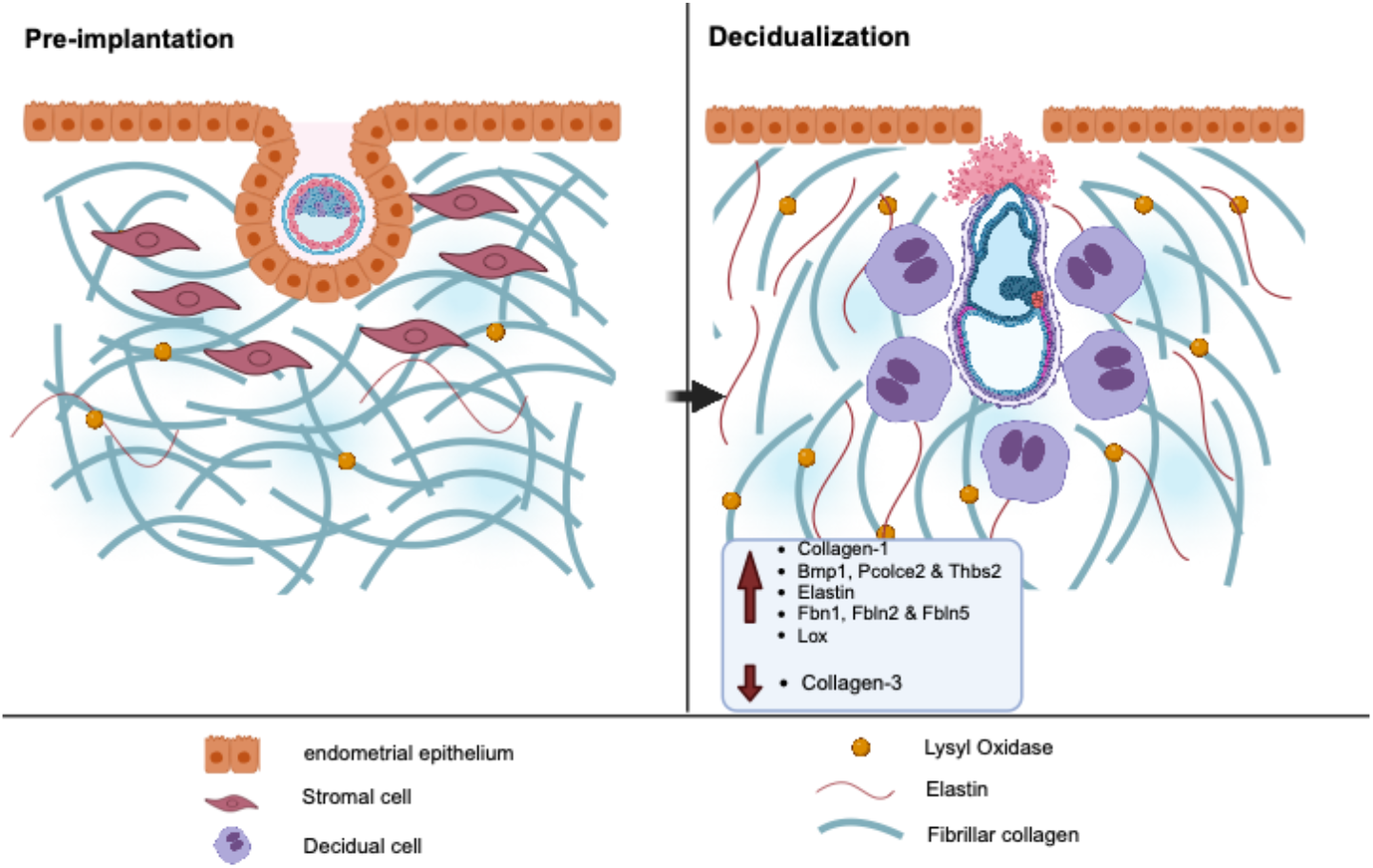

## INTRODUCTION

Embryo implantation, a pivotal step in establishing a successful pregnancy, is a multifaceted process characterized by a harmonious interplay of cellular and molecular events [1-3]. During implantation, the embryo adheres to the uterine epithelium and invades into the uterine stromal compartment [4]. This process initiates extensive proliferation and differentiation of stromal cells, leading to the formation of a transient structure called the “decidua” [5, 6]. The decidua has remarkable capabilities to receive, host, and nurture the embryo until they establish a more sophisticated communication channel through placentation [7]. Throughout this period, the decidua must maintain its functional homeostasis and structural integrity while simultaneously regulating the invasion of trophoblast cells. Decidua achieves this in part through the synthesis and reorganization of its extracellular matrix (ECM).

The tissue-specific organization of the ECM is meticulously orchestrated during embryonic development. In adulthood, the ECM undergoes limited reorganization, with the exception of reproductive tissues [8-10]. ECM reorganization, the continuous and regulated process of degrading and synthesizing new components, is pivotal for tissue development, repair, and homeostasis [11]. This reorganization is one of the fundamental events underlying decidualization triggered by embryo invasion [12-15]. Aberrant alterations in the structure and composition of the ECM in the endometrium have been associated with pathological conditions such as recurrent miscarriage, endometriosis, adenomyosis, and endometrial aging [12, 16-20]. Understanding the physiological reorganization process is essential for effectively managing disorders caused by abnormal ECM reorganization.

ECM is a sophisticated and dynamic network of proteins and polysaccharides that offers structural and biochemical support to surrounding cells [21-23]. It plays a crucial role in maintaining tissue integrity, facilitating cellular communication, and regulating various physiological processes [24, 25]. The ECM comprises a diverse array of macromolecules, including collagens, elastin, glycoproteins, proteoglycans, and glycosaminoglycans [21, 26]. Fibrillar collagens, the most abundant ECM proteins, provide tensile strength and structural support [27]. Elastin offers tissue resilience, enabling them to restore their original shape following stretching or contraction [28]. Lysyl oxidase family of enzymes play a crucial role in determining the strength of collagen and elastic fibers by catalyzing covalent crosslinking reactions [29]. Some of these factors are reported to be expressed in mouse decidua [30, 31]. However, the precise organization of multiple ECM components within the decidua, along with their spatial distribution, remains less well understood.

The study offers a detailed examination of the expression, spatial distribution, and reorganization of fibrillar collagens, elastic fibers, and lysyl oxidases within the decidua during embryo implantation. Moreover, it presents compelling experimental evidence suggesting that the endometrial stromal cells serve as the primary source of these proteins during decidualization.

## MATERIALS AND METHODS

### Mice and tissue collection

C57BL/6J129Sv mice were used in this study. To collect timed-pregnant tissues, breeding pairs were mated in the morning, and vaginal plugs were checked in the afternoon. The presence of a vaginal plug was designated as day 0 of pregnancy. Pregnant uteri were collected from gestation days 3 to 9. Implantation sites were harvested and prepared as frozen tissue blocks by immersing them in OCT (Tissue-Tek) and solidifying them with liquid nitrogen vapor. These blocks were stored at −80°C until processing. All animal experiments were conducted in accordance with the National Institutes of Health Guide for the Care and Use of Laboratory Animals humane animal care standards and were approved by the Institutional Animal Care and Use Committee of the University of Vermont.

### Second harmonic generation imaging

Frozen uterine cross sections of 50 μm in thickness were utilized in this experiment. The thawed sections were subsequently covered with 0.1 M phosphate-buffered saline (PBS) to preserve the hydration of the tissue sections during imaging. A Zeiss LSM7 inverted microscope equipped with an Achroplan 20x/0.8 objective lens was employed to acquire images of the slides. A Chameleon XR pulsed Ti: sapphire laser (Coherent, California) was adjusted to emit a wavelength of 900 nm onto the tissue, resulting in a signal detected at 450 nm. The acquired images were subsequently analyzed using ImageJ software.

### Confocal microscopic imaging

Frozen uterine cross sections (5 µm thick) were fixed in acetone for 10 minutes and subsequently blocked with 10% normal goat serum (Life Technologies) for 30 minutes at room temperature. Primary antibodies – COL1A1 (Cell Signaling, 720236S, 1:200), collagen 3 (Proteintech, 22734-1-AP, 1:500), elastin (Elastin Products Company, PR385, 1:250), LOXL1 (Thermo Fisher Scientific, PA5-87701, 1:500), LOXL2 (Abcam, Ab96233, 1:100), LOXL3 (Thermo Fisher Scientific, PA5-48462, 1:500), LOXL4 (Thermo Fisher, PA5-115520, 1:100), and smooth muscle actin (Santa Cruz, sc-32251, 1:1000) – were diluted in blocking solution, added to the tissue sections. The slides were then incubated overnight at 4°C. The following day, slides were washed with PBS and then incubated with secondary antibodies – anti-rabbit Alexa Fluor 555 (Thermo Fisher Scientific, A32732, 1:500) and anti-mouse Alexa Fluor 488 (Thermo Fisher Scientific, A11029, 1:500) – in blocking solution for 30 minutes at room temperature. Slides were washed with PBS and mounted with ProLong Gold Antifade Mountant with DNA Stain DAPI (Thermo Fisher Scientific). Images were acquired using a Nikon A1R confocal microscope equipped with a galvanometer scanner and illumination point scan at a rate of eight frames per second for a 1024 x 1024-pixel field.

### Mouse endometrial stromal cell culture

On gestation day 3, the pregnant uteri were collected, sectioned longitudinally, and subsequently cut into 3-5 mm pieces. These tissues were subsequently digested with a mixture of 6 g/L Dispase (Stem cell technologies, 07923), 25 g/L pancreatin, and antibiotic-antimycotic solution (Gibco) in HBSS for 1 hour at room temperature, followed by 15 minutes at 37°C. After aspirating the enzyme mixture, the tissues were then digested with 250 µg/mL Liberase in HBSS for 45 minutes at 37°C. The enzyme action was terminated by adding 10% FBS. The mixture was filtered through a 70 µm strainer and subsequently centrifuged. The resulting pellet was diluted in DMEM-F12 containing 2% FBS, 1% antibiotic and antimycotic solution, 1 µM progesterone, and 10 nM 17β-estradiol. Cells were counted using a hemocytometer and were plated in 6-well plates with a concentration of 1 × 10^6^ cells/well. After 2-3 hours, the culture media was replaced, and subsequently, the culture media was replaced daily. Cell lysate and conditioned media were collected at 24, 48, and 96 hours for gene and protein expression analysis.

### Quantitative polymerase chain reaction

Total RNA was extracted from stromal cell lysates using the RNeasy Mini Kit (Qiagen, 74104) following the manufacturer’s protocol. cDNA was synthesized using the iScript Reverse Transcription Supermix (Bio-Rad Laboratories). Quantitative PCR was performed using SYBR Green and primers specifically designed for the genes of interest. Gene expression was quantified using the 2^−ΔΔCt^ method, with target gene expression normalized to the housekeeping gene *Rplp0*. The primers used in this study are listed in Table 1.

**Table 1.**
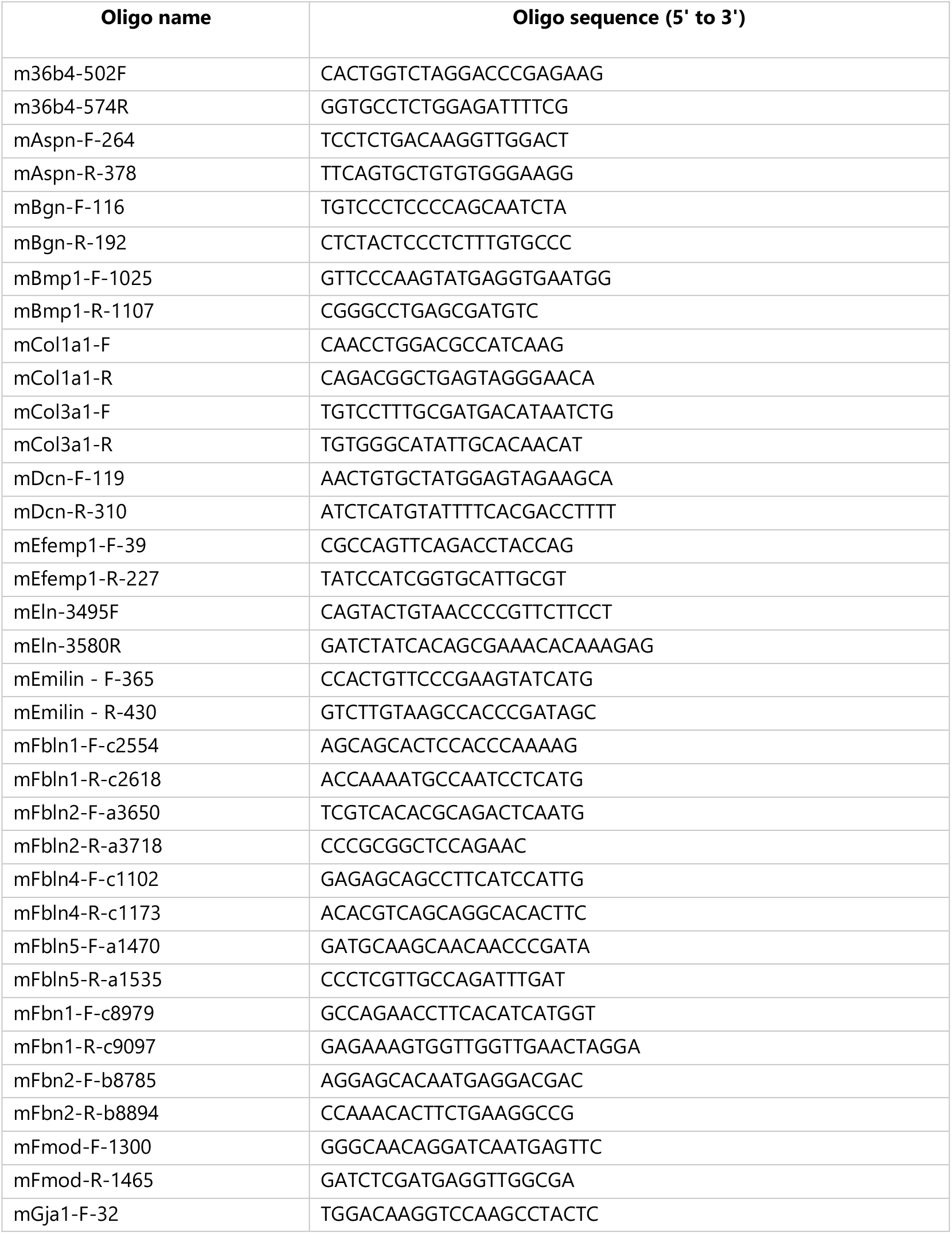
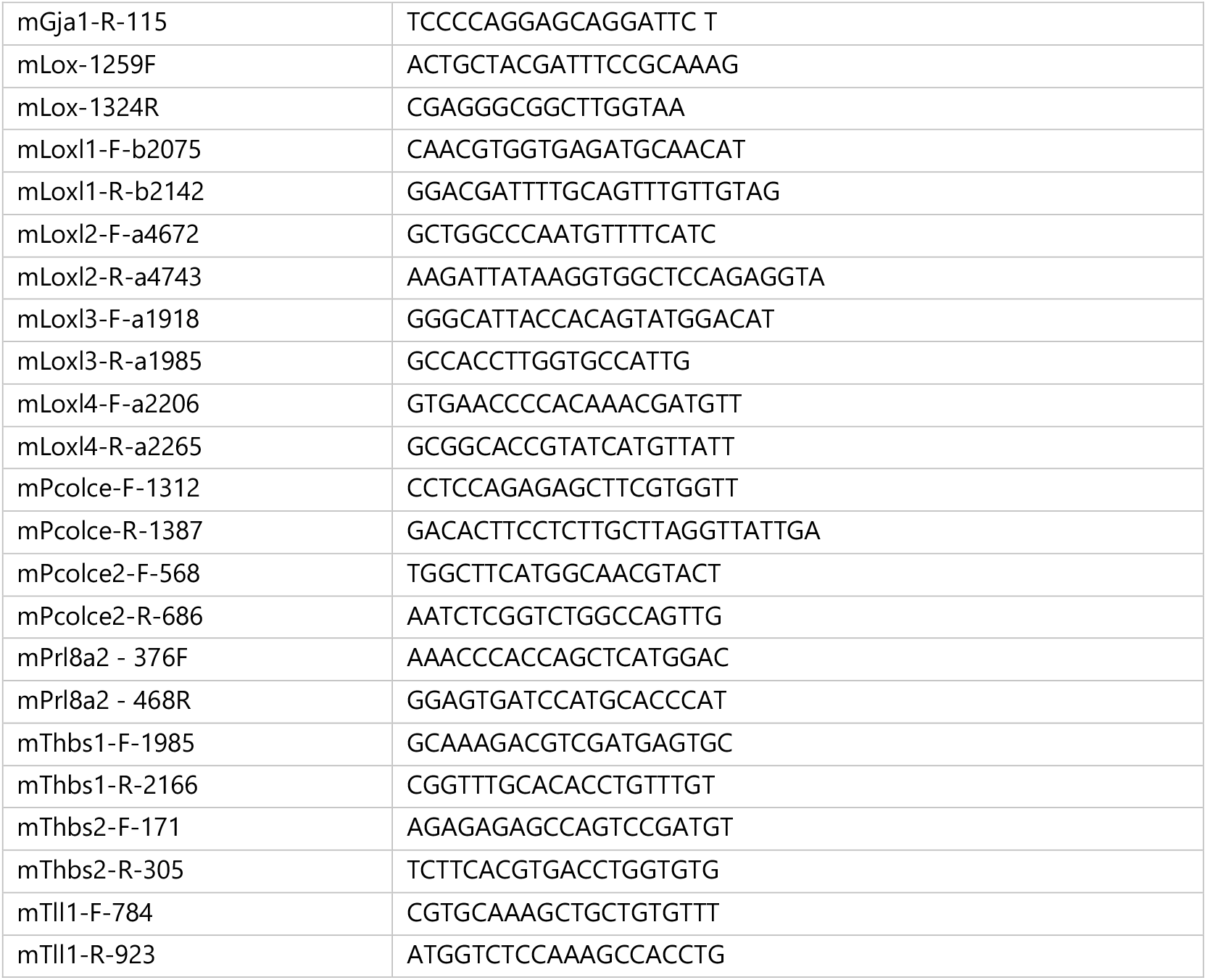
The list of primers used in this study.

### Western blot

Endometrial stromal cells were lysed using RIPA buffer containing 1% protease and phosphatase inhibitors (Thermo Fisher Scientific) and stored at −80°C until processing. Protein concentration was determined using a BCA protein assay (Thermo Fisher Scientific). Samples (20 µg) were boiled at 95°C for 10 minutes in Laemmli Sample Buffer (Bio-Rad) with β-mercaptoethanol. The samples and a protein standard (Precision Plus Protein Kaleidoscope, Bio-Rad) were loaded into a 10% Tris-HCl gel and separated at 50 V for 10 minutes, followed by 100 V for 1 hour. Proteins were then transferred onto a nitrocellulose membrane (Bio-Rad) at 100 V for 1 hour at 4°C. Membranes were blocked with 3% blotting-grade non-fat dry milk in TBST (Bio-Rad) for 1 hour at room temperature. Primary antibodies used were: COL1A1 (Cell Signaling, 72026, 1:1000), collagen 3 (Proteintech, 22734-1-AP, 1:1000), LOX (Abcam, Ab174316, 1:1000), LOXL1 (Thermo Fisher Scientific, PA5-87701, 1:500), LOXL2 (Novus, NBP1-32954, 1:500), LOXL3 (Thermo Fisher Scientific, PA5-48462, 1:1000), LOXL4 (Thermo Fisher Scientific, PA5-115520, 1:500), and GAPDH (Cell Signaling, 97166, 1:500). All primary antibodies were incubated overnight at 4°C in blocking solution. Secondary antibodies labeled with horseradish peroxidase (HRP) (Goat anti-rabbit IgG (H/L): HRP, Cell Signaling, 7074S, 1:1000; Goat anti-mouse IgG (H/L): HRP, Cell Signaling, 7076S, 1:1000) were added for 1 hour at room temperature. Imaging was conducted using Amersham ECL Western Blotting Detection Reagents and ImageQuant 800 Western blot imaging systems (Cytiva Life Sciences).

### Statistical analysis

Data were collected and analyzed using Prism software (GraphPad Software). A one-way ANOVA followed by Tukey’s multiple comparisons test was used to compare multiple groups. Values are presented as the mean ± standard error of the mean (SEM), with statistical significance determined at p<0.05.

## RESULTS

### Fibrillar collagen reorganization within the mouse decidua during embryo implantation

Second harmonic generation (SHG) imaging is a powerful tool that spatially resolves the fibrillar collagen morphology in tissues [9, 32]. Therefore, we utilized SHG imaging to examine the collagen fiber morphology during endometrial decidualization. To visualize the collagen fiber network deeper within the tissues, we acquired Z stack images at 5µm intervals through 50µm thick uterine cross sections. On gestation day 3 of the preimplantation uterus, the collagen fibers are discernible as thin strands that are evenly distributed throughout the stromal compartment. In contrast, we discovered a remarkably dense network of fibrillar collagen surrounding the invading embryo within the decidua from gestation day 6 to 8 **(Fig. 1)**. The collagen fibers oriented themselves in parallel to each other but also parallel to the invading embryo, implying that these fibers potentially determine the direction of embryo invasion from mesometrial-to-anti-mesometrial end within the decidua on gestation day 7 and 8 **(Fig. 1)**. SHG imaging revealed distinct sources and characteristics of fibrillar collagen between the mesometrial and anti-mesometrial regions. In the anti-mesometrial region, the fibrillar collagens exhibited a filament-like structure due to signals from the decidual ECM. Conversely, in the mesometrial region, they appeared tubular with branching due to signals primarily elicited by the vascular ECM on gestation days 7 and 8 **(Fig. 1)**. These findings elucidate that fibrillar collagen reorganization constitutes a distinct underlying process during decidualization.

**Fig. 1.**
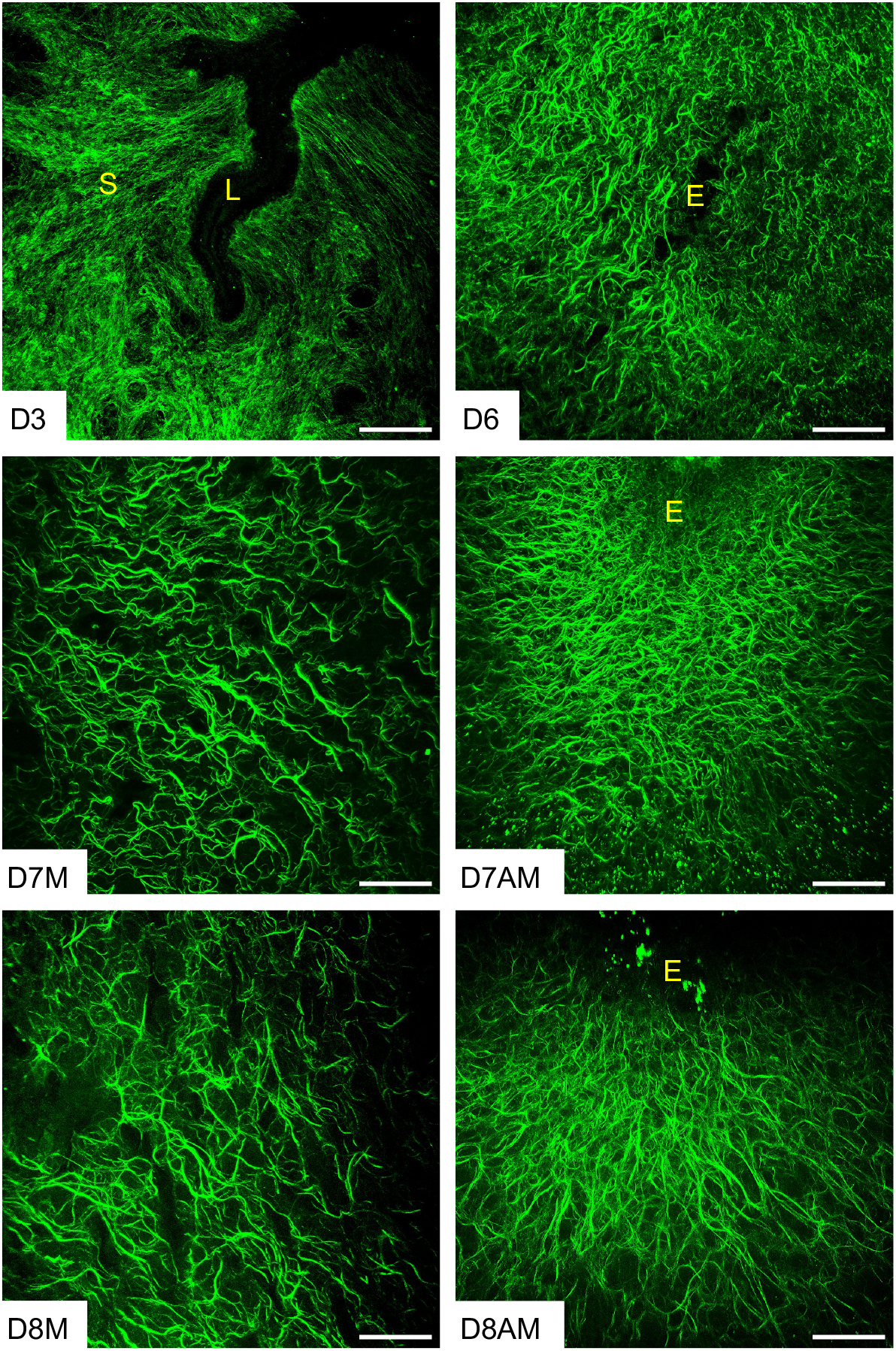
Structural reorganization of fibrillar collagen in the preimplantation and decidualized mouse uterus. Second harmonic generation imaging of collagen fibers in the gestational day 3 (D3), day (D6), day 7 [mesometrial region (D7M)/anti-mesometrial region (D7AM)], and day 8 [mesometrial region (D8M)/anti-mesometrial region (D8AM)] uteri (n=3/gestational timepoints). The Z-stack images taken at 5μm intervals deeper into 50μm thick frozen sections were projected using ImageJ. Auto fluorescent signals elicited by collagen fibers are pseudo colored in green. The imaging settings were adjusted for each individual image to optimize the visualization of morphology. L – Uterine lumen, S – Stroma, and E – Embryo. Scale bar: 100μm.

### Localization and spatial distribution of fibrillar collagens 1 and 3 in the mouse decidua

The visualization of the reorganization of fibrillar collagen by SHG led to the subsequent step of identifying the localization and spatial distribution of the primary fibrillar collagen subtypes in the decidua. To comprehend the localization and spatial distribution of the predominant fibrillar collagens, such as collagen 1 and 3, over the course of embryo implantation and decidualization, confocal imaging was employed. Consistent with SHG imaging, collagen 1 was abundantly and evenly distributed throughout the decidua. Furthermore, the collagen 1 staining exhibited a reorganization pattern surrounding the embryo **(Fig. 2)**. In contrast, collagen 3 was distributed evenly in the endometrium of gestation day 3 pregnancy **(Fig. 3)**. Notably, from gestation day 5 onwards, the spatial distribution of collagen 3 was drastically reduced within the decidua. The spatial distribution gradually faded away from the decidua but persisted surrounding the embryo and in the deep stromal compartment **(Fig. 3)**. These results demonstrate that fibrillar collagen 1 is the predominant type of collagen present within the decidua and undergoes distinct reorganization during decidualization.

**Fig. 2.**
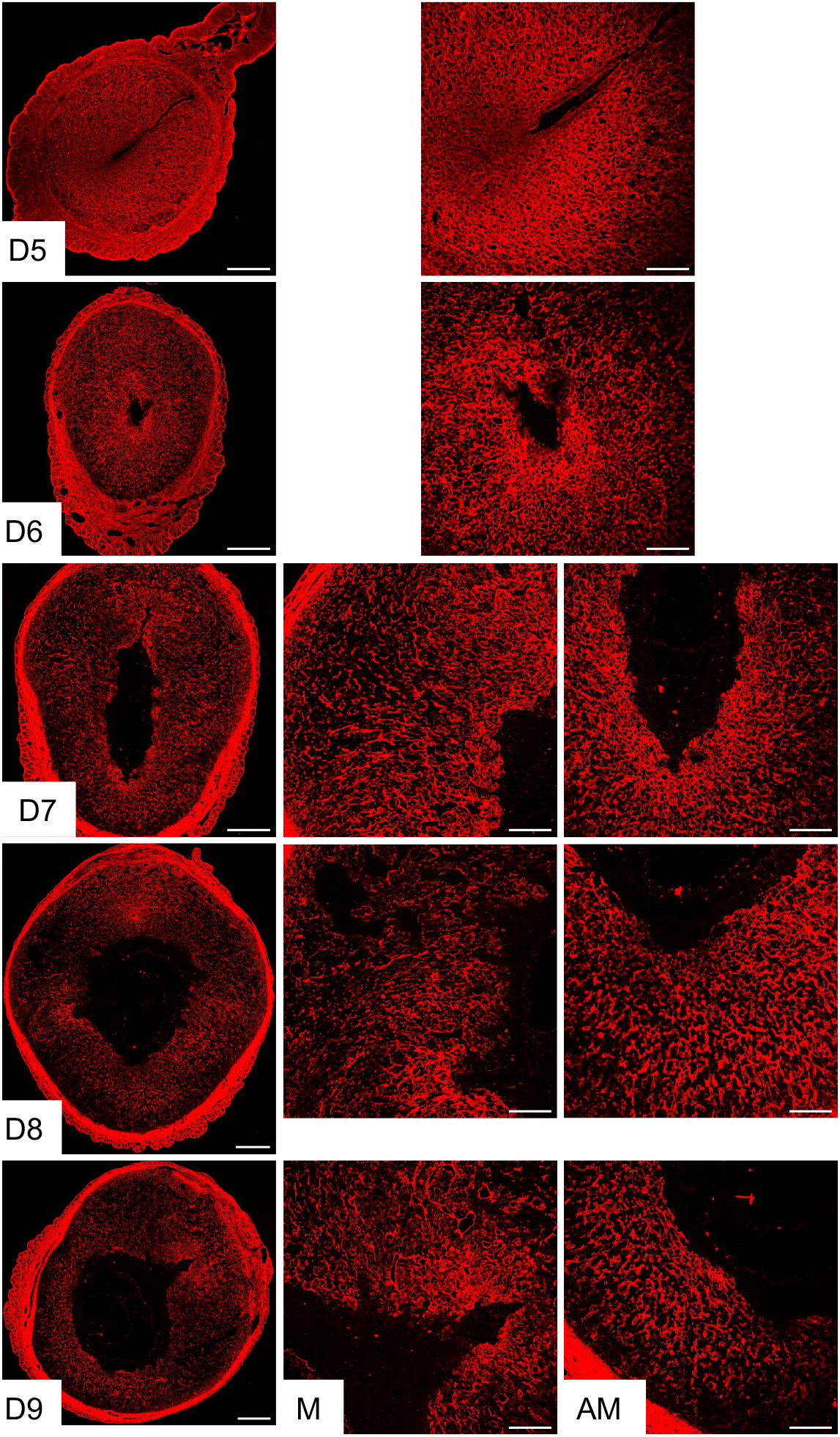
Localization of collagen 1 in the mouse endometrium during embryo implantation. Confocal imaging of COL1A1 in the frozen mouse uterine cross sections from gestation day 5 through 9. Left panel: Panoramic view of whole uterine sections; AM – Images captured from anti-mesometrial region; M - Images captured from mesometrial region. Representative images from three independent replicates. Scale bar: 500μm (left panel) and 200μm (all other images).

**Fig. 3.**
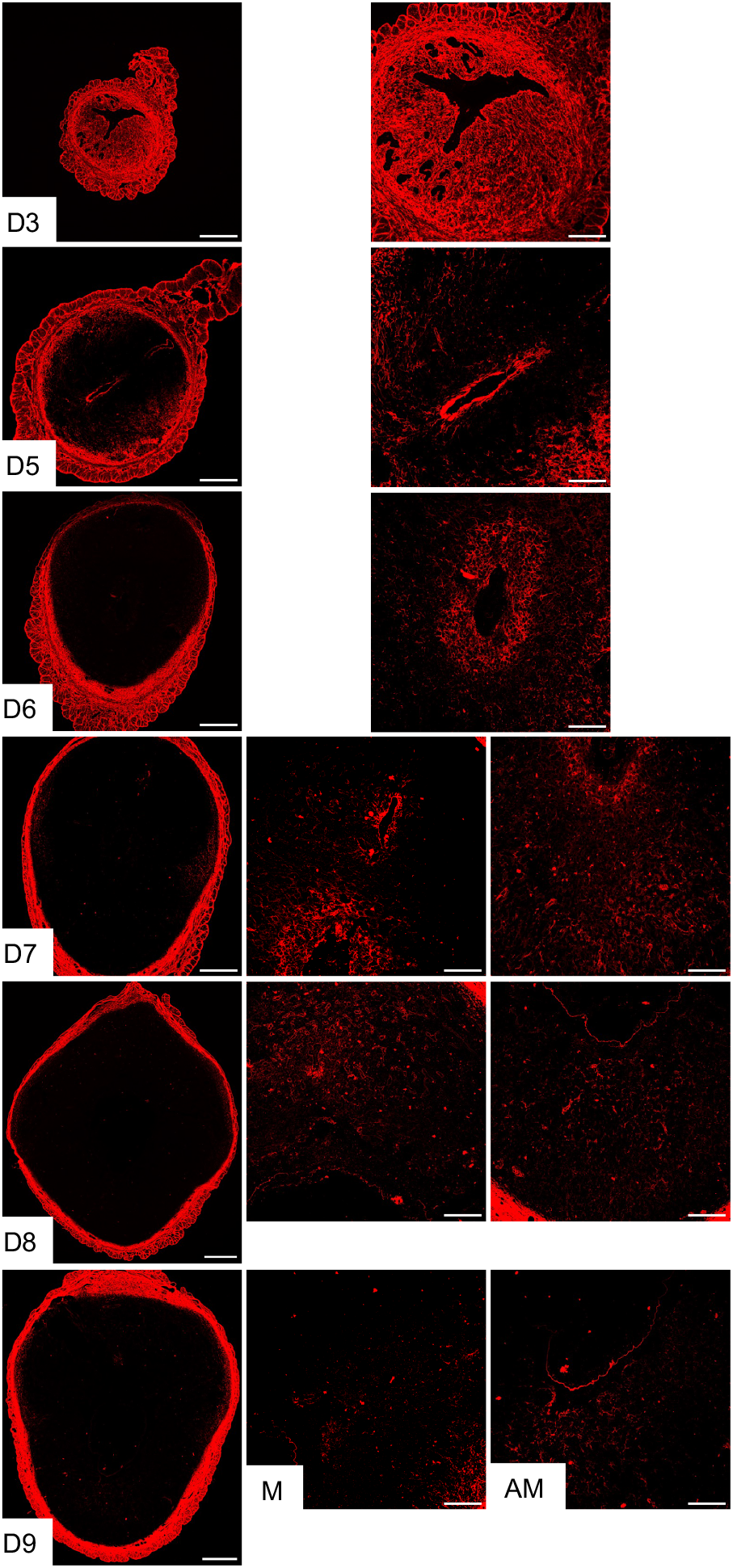
Localization of collagen 3 in the mouse endometrium during embryo implantation. Confocal imaging of collagen 3 in the frozen mouse uterine cross sections from gestation day 3, and 5 through 9. Left panel: Panoramic view of whole uterine sections; AM – Images captured from anti-mesometrial region; M - Images captured from mesometrial region. Representative images from three independent replicates. Scale bar: 500μm (left panel) and 200μm (all other images).

### Synthesis of fibrillar collagen by endometrial stromal cells *in vitro*

Our understanding of fibrillar collagen reorganization in the decidua prompted us to identify the primary source of collagen synthesis. We hypothesize that the endometrial stromal cells are the dominant source of fibrillar collagens within the decidua. To test this hypothesis, we decidualized endometrial stromal cells *in vitro* and analyzed their gene and protein expression. We collected and purified endometrial stromal cells from gestation day 3 uteri and cultured them *in vitro*. At 24, 48, 72, and 96 hours after culture, we collected cell lysates to analyze gene and protein expression. Additionally, we collected conditioned media to estimate the levels of collagen 1 and 3, as these are secreted factors. To validate our decidualization system, we analyzed the gene expression of well-known markers of decidualization, such as *Prl8a2* and *Gja1*. We observed a gradual increase in the gene expression of *Prl8a2* and *Gja1* from 24 to 96 hours after culture **(Fig. 4A)**.

**Fig. 4.**
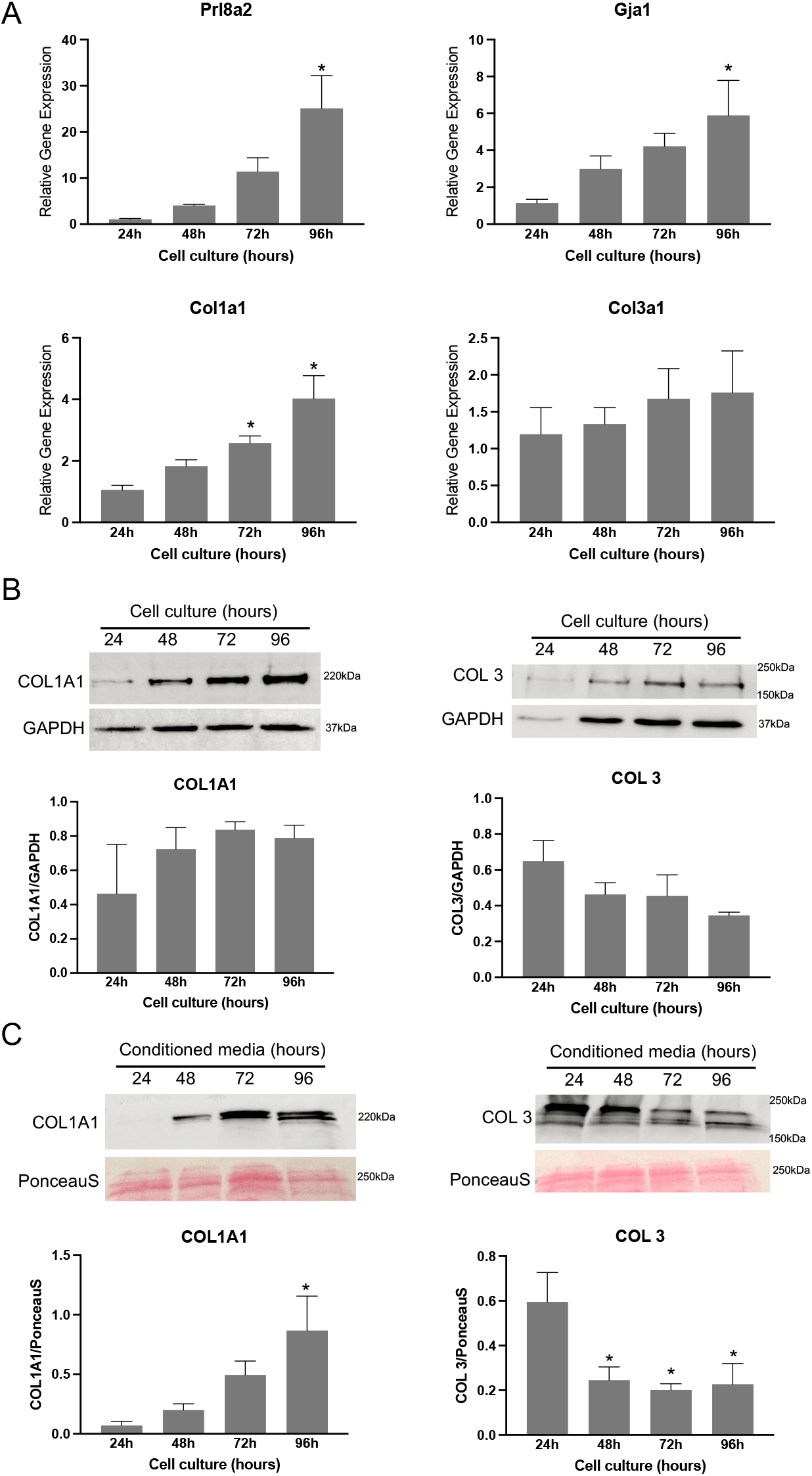
Gene expression and protein levels of fibrillar collagen 1 and 3 in mouse endometrial stromal cells decidualized *in vitro*. **A**. Gene expression of *Prl8a2* and *Gja1* and genes encoding collagen subunits, *Col1a1* and *Col3a1* in the mouse endometrial stromal cells decidualized *in vitro*. Mouse endometrial stromal cells isolated from gestation day 3 uteri were cultured *in vitro*. The cell lysates were collected at 24, 48, 72 and 96 hours to extract total RNA for gene expression analysis. The gene expression was normalized to *Rplp0* and compared with 24 hours samples (n=6/group, *p<0.05). **B**. Western blot analysis of COL1A1 and collagen 3 protein levels in cell lysates collected at 24, 48, 72 and 96 hours. GAPDH was used as a loading control. These are representative images from three independent replicates. Quantification of protein levels of COL1A1 and collagen 3 is shown in histogram. **C**. Western blot analysis of COL1A1 and collagen 3 protein levels in the conditioned media collected at 24, 48, 72 and 96 hours after culture. These are representative images from four independent replicates. Note: The culture medium was replaced every day. PonceauS stain was used to assess transfer efficiency and to visualize total proteins. Quantification of protein levels of COL1A1 and collagen 3 is shown in histogram (n=4/group, *p<0.05).

The gene expression of *Col1a1* gradually increased and reached a significant level at 72 hours, with a maximum at 96 hours, compared to 24 hours. On the other hand, the gene expression of *Col3a1* remained relatively constant throughout the time points tested **(Fig. 4A)**. We were able to identify the COL1A1 and collagen 3 proteins in the cell lysates, but their levels did not show significant differences **(Fig. 4B)**. In contrast, the protein levels of COL1A1 exhibited a steady increase in the conditioned media collected and reached a statistically significant level at 96 hours. Conversely, the levels of collagen 3 were significantly reduced at all time points compared to 24 hours **(Fig. 4C)**. These results correlate with localization studies described above for collagen 1 and 3 (Fig. 2 and 3). These findings collectively demonstrate that endometrial decidual cells are the primary source of fibrillar collagen during decidualization in response to embryo implantation.

### Expression of genes encoding factors involved in collagen synthesis, processing, and assembly during *in vitro* decidualization

To further understand the process of fibrillar collagen reorganization, we examined the expression of genes involved in its synthesis, processing, and assembly. Procollagen molecules, the soluble precursors of collagen, consist of globular pro-peptides at their N and C ends. The proteolytic action of procollagen N and C proteinases converts procollagen into mature collagen. These proteinases encompass bone morphogenetic protein 1 (BMP1), tolloid-like families (Tll), and a disintegrin and metalloproteinase with thrombospondin motif (ADAMTS). The activity of procollagen proteinases is regulated by enhancers, such as procollagen C-endopeptidase enhancer-1 (Pcolce) and procollagen C-C-endopeptidase enhancer-2 (Pcolce2) [27, 33]. Notably, the expression of *Bmp1, Pcolce*, and *Pcolce2* gradually increases over time during stromal cell decidualization *in vitro*, reaching a peak significant level at 96 hours, while *Tll1* levels remain constant throughout **(Fig. 5A)**.

**Fig. 5.**
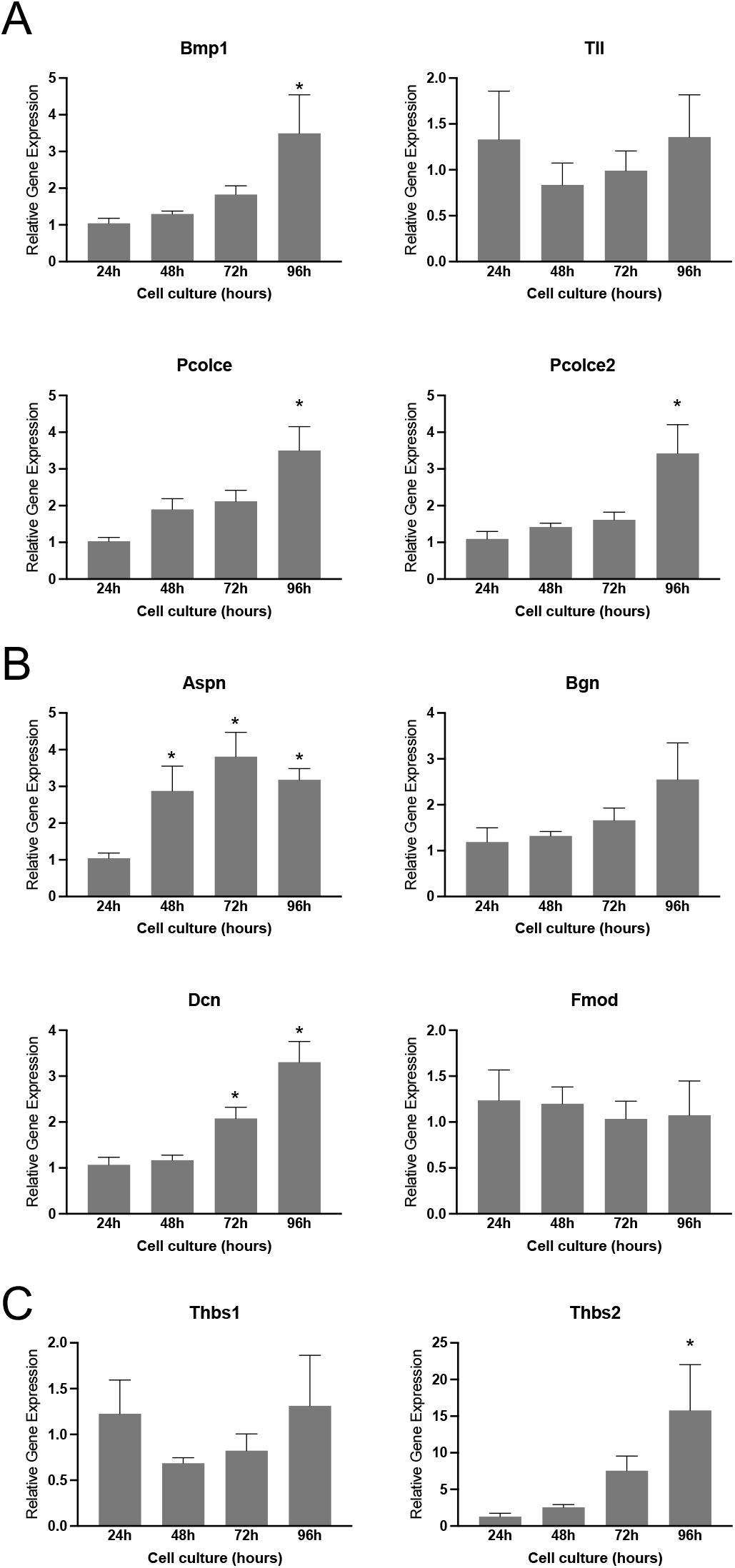
Gene expression of factors involved in the synthesis, processing, and assembly of fibrillar collagen in mouse endometrial stromal cells decidualized *in vitro*. **A**. Gene expression of *Bmp1, Tll, Pcolce*, and *Pcolce2*, in the mouse endometrial stromal cells decidualized *in vitro*. **B**. Gene expression of genes encoding small leucine rich proteoglycans such as *Aspn, Bgn, Dcn*, and *Fmod* in the mouse endometrial stromal cells decidualized *in vitro*. **C**. Gene expression of *Thbs1* and *2* in the mouse endometrial stromal cells decidualized *in vitro*. Mouse endometrial stromal cells isolated from gestation day 3 uteri were cultured *in vitro*. The cell lysates were collected at 24, 48, 72 and 96 hours to extract total RNA for gene expression analysis. The gene expression was normalized to *Rplp0* and compared with 24 hours samples (n=6/group, *p<0.05).

Small leucine-rich proteoglycans (SLRPs), including asporin, biglycan, decorin, fibromodulin, and lumican, play a crucial role in collagen size, content, morphology, and growth rate during microfibril growth and merging into fibrils in both the longitudinal and axial directions [34, 35]. The gene expression of these SLRPs reveals that asporin and decorin exhibit significant induction during decidualization compared to other factors **(Fig. 5B)**. Thrombospondin-1 (*Thbs1*) and thrombospondin-2 (*Thbs2*) have both direct and indirect effects on collagen homeostasis [36, 37]. Notably, the expression of *Thbs2* is significantly elevated at 96 hours, while *Thbs1* levels remain unchanged **(Fig. 5C)**. These findings suggest that the genes encoding factors influencing collagen synthesis, processing, and assembly exhibit differential regulation within the decidua, indicating alterations in the synthesis and processing of fibrillar collagen during endometrial decidualization.

### Elastic fiber reorganization within the mouse decidua during embryo implantation

Elastic fibers, in conjunction with collagen, constitute the fibrous components of the ECM and are an integral component of the vasculature, conferring tissues with elasticity and resilience [38, 39]. These properties render elastic fibers not only indispensable for the integrity of the decidua but also essential for decidual angiogenesis. The decidua undergoes transformation into the maternal placenta during placentation, which is predominantly composed of an intricate network of vasculature [40]. Therefore, we focused on the reorganization of elastic fibers within the decidua. Elastic fibers are composed of cross-linked tropoelastin embedded on microfibrils. Microfibrils are primarily made up of fibrillin and contain other proteins like EMILIN-1, fibulins, and microfibril-associated glycoproteins [38]. To visualize the reorganization of elastic fibers in the decidua, we used confocal imaging to detect elastin. Our observations showed that elastin expression began to appear around the implanted embryo within the decidua on gestational day 5 and gradually increased to cover the entire decidua by gestational day 8 **(Fig. 6)**. Notably, on gestational day 9, the elastin was heavily localized in the blood vessels located within the mesometrial region, which will later form the placenta.

**Fig. 6.**
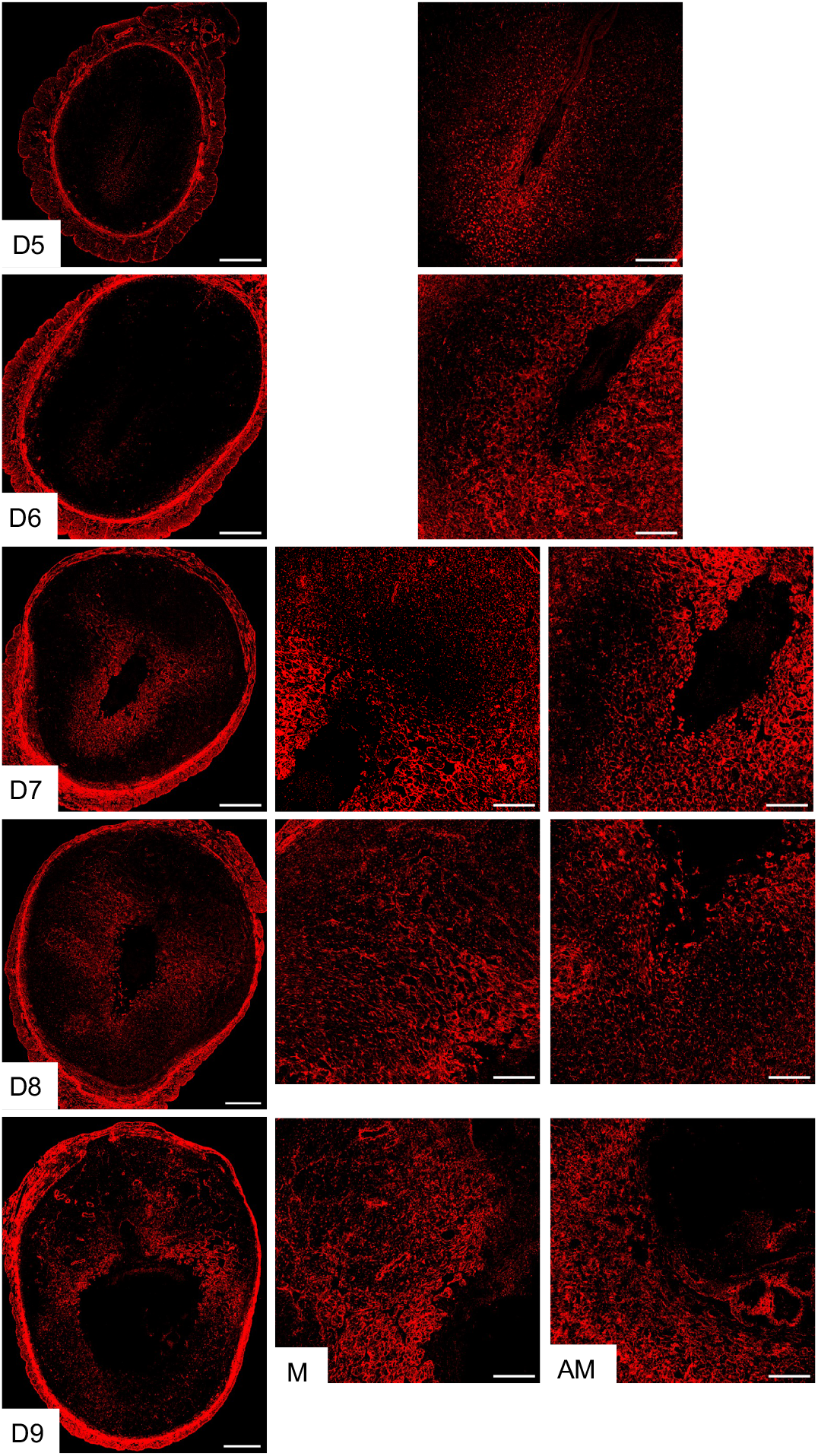
Localization of elastin in the mouse endometrium during early pregnancy. Confocal imaging of elastin in the frozen mouse uterine cross sections from gestation day 5 through 9. Left panel: Panoramic view of whole uterine sections; AM – Images captured from anti-mesometrial region; M - Images captured from mesometrial region. Representative images from three independent replicates. The imaging settings were adjusted for each individual image to optimize the visualization of morphology. Scale bar: 500μm (left panel) and 200μm (all other images). gestation day 7, then became widespread throughout the decidua on gestation day 8 **(Fig. 9)**. These findings demonstrate that all lysyl oxidases are expressed in the endometrium during decidualization.

Subsequently, we investigated the role of endometrial stromal cells in the production of elastic fibers. To achieve this objective, we conducted a comprehensive analysis of gene expression related to elastin synthesis, processing, and assembly in cell lysates obtained from endometrial stromal cell culture. While the expression of elastin, fibrillin 2, emilin, fibulin 1, and 3 was detected, no significant difference was observed during endometrial decidualization *in vitro*. However, the genes such as fibrillin1 and fibulin 2, 4, and 5 were induced significantly at 96 hours **(Fig. 7)**. These findings suggest that the reorganization of elastic fibers in the endometrium during decidualization may assume a pivotal role in the process of placentation.

**Fig. 7.**
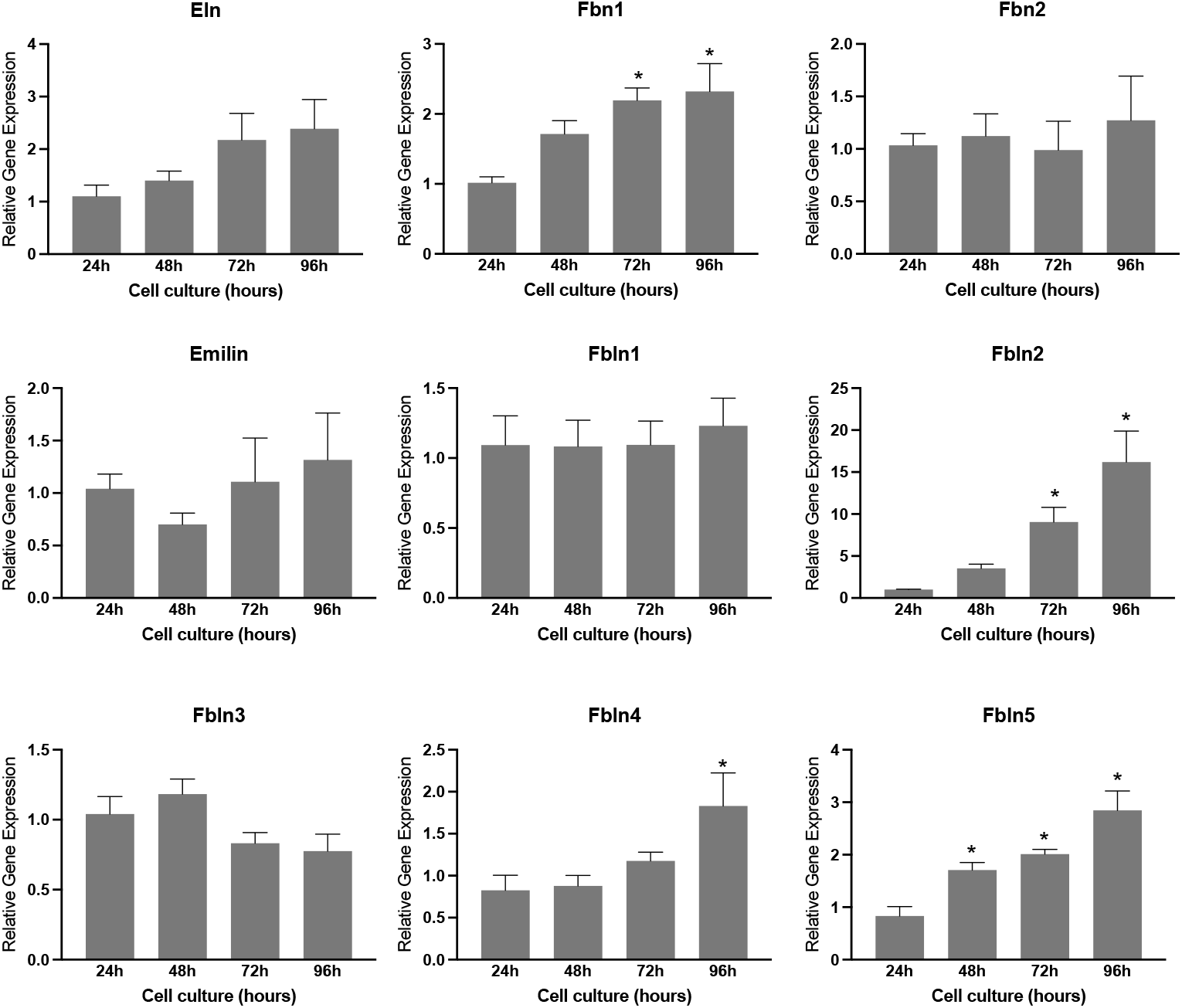
Gene expression of factors involved in the synthesis, processing, and assembly of elastic fiber in in mouse endometrial stromal cells decidualized *in vitro*. Gene expression of *Eln, Fbn1, Fbn2, Emillin, Fbln1, Fbln2, Fbln3m Fbln4*, and *Fbln5* in the mouse endometrial stromal cells decidualized *in vitro*. Mouse endometrial stromal cells isolated from gestation day 3 uteri were cultured *in vitro*. The cell lysates were collected at 24, 48, 72 and 96 hours to extract total RNA for gene expression analysis. The gene expression was normalized to *Rplp0* and compared with 24 hours samples (n=6/group, *p<0.05).

### Expression and localization of lysyl oxidases in the decidua during embryo implantation

Collagen and elastic fibrous proteins are primarily strengthened by covalent crosslinking of lysine residues, catalyzed by the lysyl oxidase family of enzymes [29, 41]. This family comprises five members: Lysyl oxidase (LOX) and LOX-like 1 to 4 (LOXL1-4). These enzymes share the uniform ability to cross-link collagen and elastic fibers, but they exhibit tissue-specific spatiotemporal expression and play multiple roles in the ECM and tissue function [42, 43]. Consequently, we investigated the expression of lysyl oxidases during decidualization. LOX has been reported to be expressed within the decidua during embryo implantation [31]. In this study, we focused on the localization of LOXL1-4 in the decidua. LOXL1 expression was modest within the decidua surrounding the implanted embryo, but it gradually expanded spatially from gestation day 7 to 8. LOXL2 was evenly distributed throughout the decidua from gestation day 5 to 8 **(Fig. 8)**. LOXL3 was primarily localized in the primary decidual zone on gestation day 5 and 6, then gradually expanded throughout the decidua from gestation 7 to 8. LOXL4 was first detected on gestation day 6 and diffusely expressed in the secondary decidual region on

**Fig. 8.**
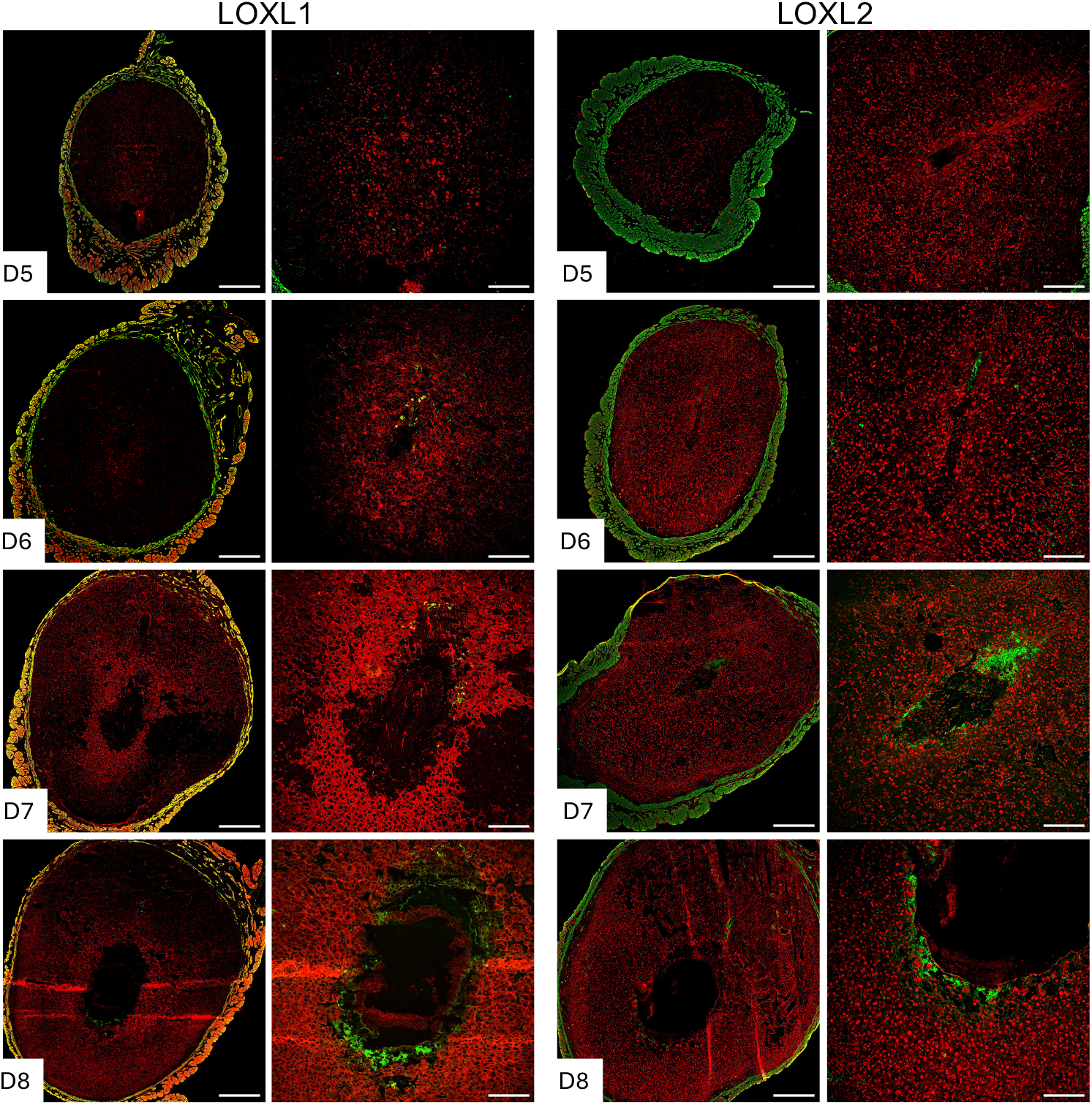
Localization of LOXL1 and LOXL2 in the mouse endometrium during embryo implantation. Confocal imaging of LOXL1 and LOXL2 in the frozen mouse uterine cross sections from gestation day 5 through 8. Left panels: Panoramic view of whole uterine sections. Red – LOXL1 or LOXL2; Green - Smooth muscle actin. Right panels: Images captured in decidua. Representative images from three independent replicates. The imaging settings were adjusted for each individual image to optimize the visualization of morphology. Scale bar: 500μm (left panel) and 200μm (all other images).

**Fig. 9.**
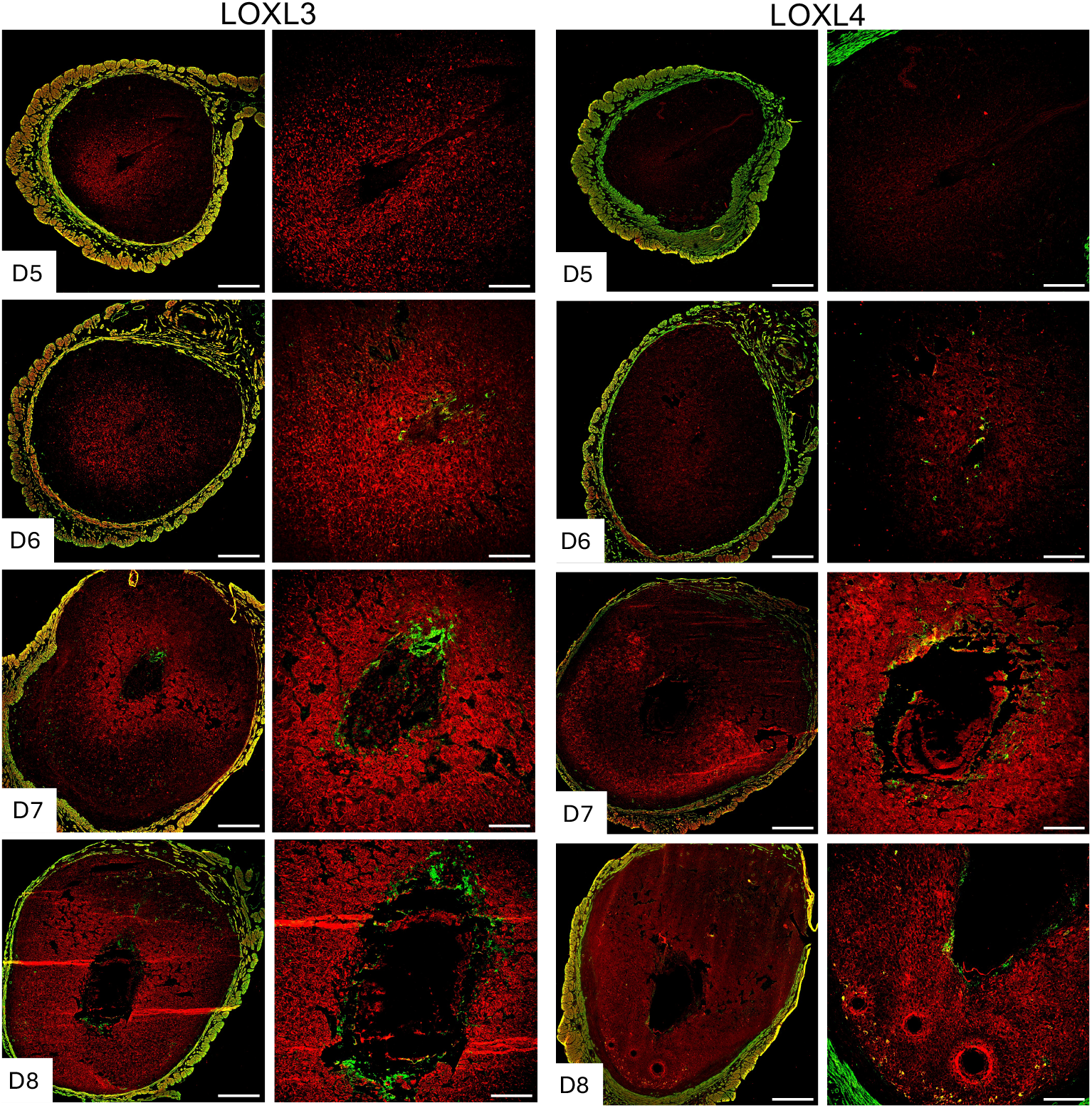
Localization of LOXL3 and LOXL4 in the mouse endometrium during embryo implantation. Confocal imaging of LOXL3 and LOXL4 in the frozen mouse uterine cross sections from gestation day 5 through 8. Left panels: Panoramic view of whole uterine sections. Red – LOXL3 or LOXL4; Green - Smooth muscle actin. Right panels: Images captured in decidua. Representative images from three independent replicates. The imaging settings were adjusted for each individual image to optimize the visualization of morphology. Scale bar: 500μm (left panel) and 200μm (all other images).

To elucidate the role of endometrial stromal cells in the synthesis of lysyl oxidases, we analyzed the gene and protein expression of lysyl oxidases in lysates derived from endometrial stromal cell cultures. Our findings revealed the expression of all five lysyl oxidase genes in decidual cells. Among all genes, *Lox* and *Loxl4* were significantly induced at 96 hours **(Fig. 10A)**. Consistent with gene expression, the protein level of LOX was significantly elevated from 48 hours to 96 hours compared to 24 hours. LOXL1, LOXL2, and LOXL3 levels remained unchanged throughout the *in vitro* decidualization period. In contrast to its gene expression, LOXL4 protein levels were significantly reduced during *in vitro* decidualization **(Fig. 10B, C)**. This discrepancy could be attributed to its post-translational processing within the ECM. These findings suggest that lysyl oxidases are synthesized by endometrial stromal cells during decidualization and might play a crucial role in modifying the structural integrity of the decidua by stabilizing collagen and elastic fibers.

**Fig. 10.**
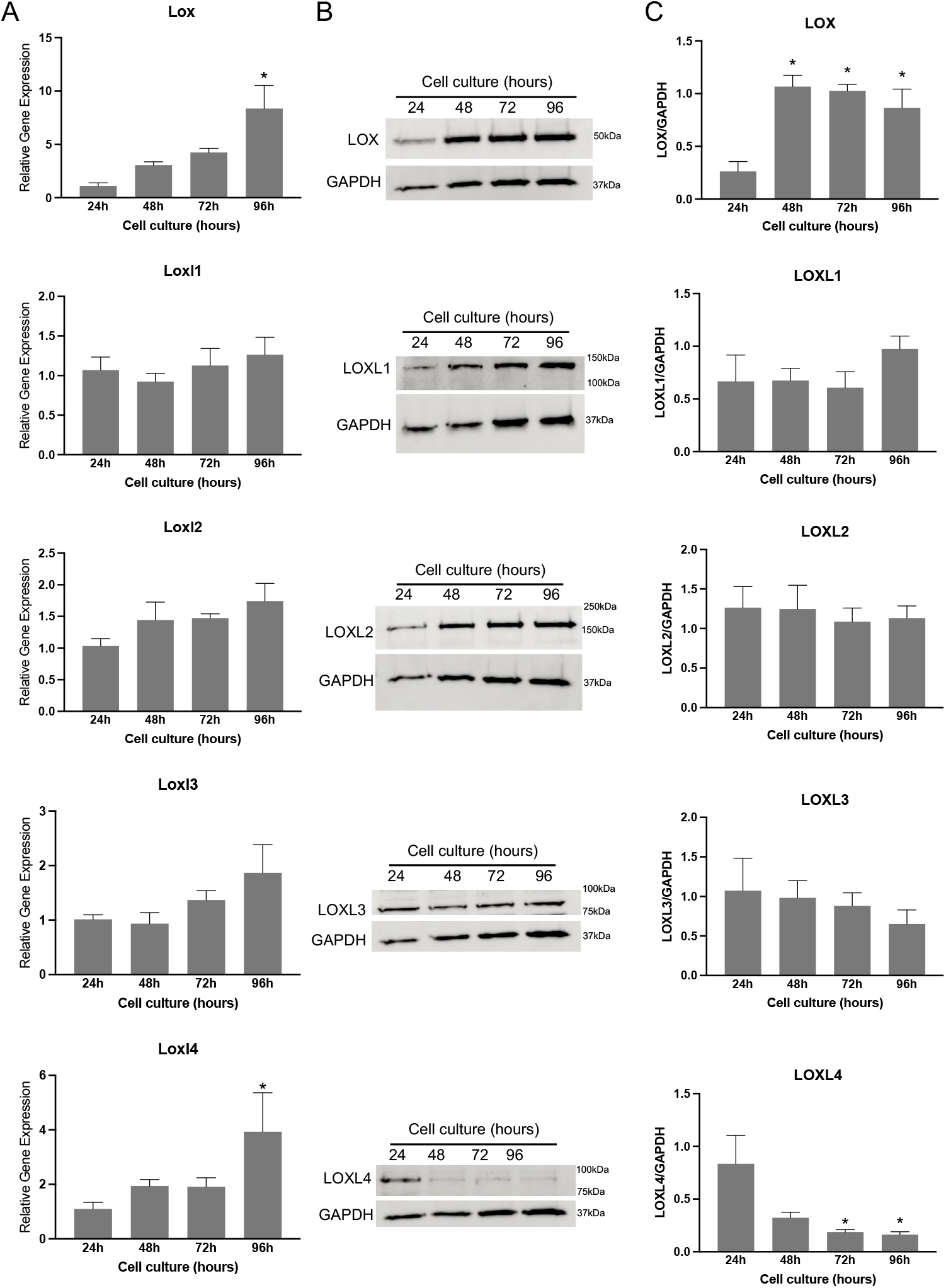
Gene expression and protein levels of LOX and LOXL1-4 in mouse endometrial stromal cells decidualized *in vitro*. **A**. Gene expression of *Lox* and *Loxl1-4* in the mouse endometrial stromal cells decidualized *in vitro*. Mouse endometrial stromal cells isolated from gestation day 3 uteri were cultured *in vitro*. The cell lysates were collected at 24, 48, 72 and 96 hours to extract total RNA for gene expression analysis. The gene expression was normalized to *Rplp0* and compared with 24 hours samples (n=6/group, *p<0.05). **B**. Western blot analysis of LOX and LOXL1-4 protein levels in cell lysates collected at 24, 48, 72 and 96 hours. GAPDH was used as a loading control. These are representative images from three independent replicates. **C**.Quantification of protein levels of LOX and LOXL1-4 is shown in histogram (n=3/group, *p<0.05).

## DISCUSSION

This study elucidates the ECM reorganization that occurs during endometrial decidualization, a pivotal phase in embryo implantation. It provides an in-depth analysis of the expression, distribution, and reorganization of fibrillar collagens, elastic fibers, and lysyl oxidases within the decidua during the implantation period. We have observed a rise in fibrillar collagen 1 levels and a decline in collagen 3 levels during the decidualization process. These findings are consistent with the reduction in the ratio of collagen type 3 to type 1 in human decidua [44, 45]. Utilizing SHG, a highly sophisticated tool capable of mapping fibrillar collagen structure deeper into tissues, we have recorded the fibrillar collagen reorganization of the decidual tissues surrounding the invading embryo over the course of embryo implantation. The remarkable alterations in fiber structure and orientation between preimplantation and decidualized uterine stroma offer a novel perspective on the process of ECM reorganization during decidualization. The attachment of the embryo to the uterine epithelium and its subsequent invasion into the uterine stromal compartment occurs in the mesometrium to anti-mesometrial orientation. This process remains an enigma for endometrial biologists. The collagen fiber orientation within the decidua during embryo invasion provides a potential clue to this process. Based on our observations obtained through SHG imaging of fibrillar collagen, we propose that the reorientation of the collagen fibers in parallel to the invading embryo may contribute to the direction of mesometrial to anti-mesometrial invasion of the embryo with the decidua.

Elastic fibers, a crucial fibrous component that collaborates with collagen to maintain the ECM architecture, are also an integral component of the vascular wall, providing tissue elasticity and resilience [28, 38]. Elastin is highly localized in the mesometrial decidua (the destined site of placentation) and its blood vessels on gestation day 9. This spatial distribution of elastic fibers within the decidua over the implantation period suggests a potential role in decidual angiogenesis and, consequently, placentation. The strength and stability of collagen and elastic fibers are attributed to their cross-linking mediated by lysyl oxidases [29, 43]. Although these lysyl oxidases are widely expressed in collagen and elastin-containing tissues, their spatio-temporal expression pattern varies across different tissues [43]. The robust spatial distribution of these lysyl oxidases within the decidua during embryo implantation implicates their potential role in embryo implantation.

The forceful invasion of trophoblast into decidual tissues has the potential to alter the structural integrity and, consequently, the mechanical properties of the decidua. Therefore, the decidua should possess appropriate mechanisms to regulate trophoblast invasion without compromising its tissue mechanical homeostasis. The embryo’s invasion and its impact on decidual tissue stiffness throughout decidualization remain poorly understood. Based on our current understanding, we propose that ECM reorganization during decidualization assumes this responsibility to maintain decidual tissue mechanical homeostasis by precisely regulating embryo invasion. This hypothesis is supported by clinical complications such as shallow implantation and placenta accreta spectrum disorders. Unregulated, deeper trophoblast invasion that penetrates the myometrium can lead to placenta accreta, while inadequate invasion results in shallow implantation [15, 46, 47]. The rapid proliferation and differentiation of stromal cells lead to the development of enlarged tissue growth at the implantation sites, known as decidua or deciduoma. Although not a true neoplasm, decidua exhibits tissue growth and ECM reorganization, similar to neoplastic processes. Notably, fibrillar collagen reorganization is closely associated with tumor tissue stiffness in neoplasm [22, 48-50]. Aberrant collagen reorganization and elevated lysyl oxidase levels has also been observed in endometrial pathologies, such as endometriosis [18, 19]. Consequently, it is imperative to expand our understanding of endometrial tissue mechanical properties during embryo implantation.

ECM is predominantly synthesized by stromal cells, although most other cell types, such as smooth muscle cells, immune cells, and vascular endothelial cells, are also known to synthesize ECM constituents [28, 51]. Utilizing purification and culture of endometrial stromal cells, we have unequivocally demonstrated that uterine stromal cells can synthesize fibrillar collagen, elastin, and lysyl oxidases during decidualization even in the absence of embryo invasion. These findings confirm that endometrial stromal cells assume a central role in orchestrating this ECM reorganization. Embryonic life customizes ECM, which undergoes limited reorganization in adulthood, except in reproductive tissues [8-10]. ECM reorganization, including its rapid degradation and turnover, has been reported in various reproductive tissues [52-54]. The timely ECM reorganization is crucial for reproductive tissues to fulfill their functions, which are contingent on the dynamic reproductive statuses of non-pregnancy and pregnancy. For instance, the mechanisms involved in ECM reorganization in the myometrium and cervix to perform opposing functions during pregnancy and parturition have been elucidated [9, 52, 55, 56]. Any aberrant ECM reorganization in these tissues can lead to parturition defects such as dystocia and preterm birth [55, 57]. Defective ECM, particularly elastic fibers and their constituents, in the pelvic floor is the predominant cause of pelvic organ prolapse [10, 58]. These studies underscore the significance of ECM reorganization in the reproductive tract. Our knowledge regarding ECM reorganization in the endometrium is still limited to the processes of decidualization, placentation, and endometrial regeneration.

The present study elucidates several clues involved in ECM reorganization during endometrial decidualization and embryo implantation. Future research endeavors within our laboratory will involve the generation of mouse models deficient in specific ECM components to ascertain their functional relevance *in vivo* during these processes.

## Acknowledgements

We sincerely acknowledge Dr. Douglas Taatjes, Director and Nicole Bouffard, staff at the Microscopy Imaging Center at the University of Vermont (RRID# SCR_018821) for their help and support. Confocal microscopy was performed on a Nikon A1R-HD point scanning confocal supported by NIH award number 1S10OD025030-01 from the Office of Research Infrastructure Programs.

## Author contributions

SN conceptualized and designed the study. SN, MG, SM, and RD conducted experiments, and analyzed data. MG wrote the first draft of the manuscript. All authors reviewed and approved the manuscript.

## Data availability

The data underlying this article are available in the article.

